# Physico-chemical and microbiological characteristics of honey produced by stingless bees *(Meliponula beccarii)* from the Oromia Region, Ethiopia

**DOI:** 10.1101/2022.12.23.521720

**Authors:** Teferi Damto, Deresa Kebeba, Meseret Gemeda

## Abstract

The study was designed to characterize the Physico-chemical and microbiological properties of stingless bee honey (Meliponula beccarii L.) in the Oromia Region, Ethiopia. About thirty-nine (39) honey samples were collected from underground soils through an excavation of natural nests. The study revealed that the mean values of physicochemical parameters for honey samples were: moisture content (30.69±0.29), ash content (0.16±0.01), electrical conductivity (0.44±0.2), pH (3.30±0.03), free acidity (92.39±4.45), HMF (6.58±0.36), fructose (36.48±0.54), glucose (27.67±0.43),sucrose(1.24±0.18), maltose (1.2±0.18) and reducing sugar (64.15±0.75). Stingless bee honey in this study is characterized as possessing higher moisture content and free acidity, but a lower level of sugar as compared to Apis mellifera honey standard. The occurrence of microorganisms in the stingless honey sample ranged from 2.55 × 10 ^4^ to 1.9 × 10^3^ CFU/ml for Aerobic Mesophilic, 1.68 x10^4^ to 9 x10^2^ CFU/ml for yeast, and 1.8 x10^3^ to 2 x10^2^ CFU/ml for mold. The number of aerobic spore-forming bacteria was at non-detectable levels in all samples while Staphylococci species was detectable only in a sample obtained from Guduru. This indicated that there might be contamination during the harvesting, processing, and storage of stingless bee honey samples. The microbiological and physicochemical properties of stingless bee honey are different from those of A. mellifera honey and need to establish for specific quality standards to promote its commercialization.

## 1. Introduction

Honey is one of the most complex natural foodstuffs; it is mainly composed of sugars and other constituents, such as enzymes, amino acids, organic acids, carotenoids, vitamins, minerals, and aromatic substances. It is rich in flavonoids and phenolic acids that exhibit a wide range of biological effects and which also act as natural antioxidants [1]. Honey composition is varied and is linked to several factors that directly affect its composition and quality such as the bee species, floral origin, and environmental and storage conditions[2].

World honey production and consumption are based on the product obtained from the species Apis mellifera. However, other small productions are based on products obtained from other species of bees, such as the stingless bee [3]. Stingless bee honey production is limited compared with *A. mellifera* honey since it does not reach industrial levels, has a lower shelf life, lacks quality standards, and has little knowledge about the product [4]. The significant difference is mainly contributed by the smaller body size of stingless bees which helps them to collect nectar from various nectar sources including small flowers that cannot be penetrated by honey bees. However, the consequences of having a smaller body are lesser production of honey annually and making the market price per kg higher. Stingless bees collect nectar from the nectar glands of flowers and other parts of plants. After collection, the nectar is transformed and stored in honey cells (pots) [5]. Different compositions of honey in each area show different tastes as well as medicinal properties depending on the foraged food sources[6,7]. In Ethiopia, both honey types are produced and commercialized all over the country.

Nowadays, stingless bee honey is gaining more attention worldwide as it is revealed to havetherapeutic properties which potentially can help in wound healing, and eye diseases, and also can pose as an anti-inflammatory agent [8]. On other hand, this honey is much appreciated in the region for fresh consumption, having a more acid taste, higher fluidity, and lower viscosity, in addition to its medicinal appeal, which increases and raises the price of the product. Stingless bees produce and store much less honey on a per hive basis (1-5 kg of honey per year depending on the species) compared to *Apis mellifera* bees, with an average of 20 kg of honey per hive[9]. For this reason, stingless bee honey is available in traditional markets and commands a significantly higher price compared to *A. mellifera*. But, the production and commercialization of honey from meliponines is still limited when compared to the *Apis* genus.

Numerous studies have been done on the physicochemical properties of stingless bee honey from around the world, such as those from Venezuela [10], Thailand [11], and Mexico[12]. The principal studies on Ethiopia honey have focused mainly on the characterizing physicochemical parameters, and botanical and geographical properties of honey from *A. mellifera* [13,14,15,16, 17,18,19,20,21]. Despite a large number of stingless honey bees produced and higher market demand, none of the studies were reported on the physicochemical parameters and microbiological properties of stingless bee honey in Ethiopia, with exception of studies by Gela *et al*. [22] focused only on physicochemical properties of stingless bee honey from West Shoa Zone. Insufficient knowledge about stingless bee honey characterization makes it difficult to establish an official standard of quality needed to expand its production and marketing. Quality indicators established in Ethiopia legislation refer to parameters for honey from the *Apis mellifera* species [23], which do not correspond to those observed in melliponine honey. Stingless bee honey is also not included in international standards for honey [5,24]. The study of its microbiological characterization and physicochemical composition is important for the certification process, which determines quality and origin (geographic and botanical), and assists in an inspection. Thus, to increase the knowledge about the shelf life quality of honey from stingless bees, this study aimed to determine the physicochemical and microbiological characteristics of *M. beccarii* honey produced in different locations in Ethiopia.

## 2. Materials and Methods

### 2.1. Study area Description

Potential districts for stingless bee honey production were purposively selected for sample collection. Then, stingless bee honey samples were collected from eight (8) zones of the Oromia Region (West Shoa, Horro Guduru Wollega, East Wollega, West Wollega, B/Bedelle, I/A/Boor, Jimma, and Guji) and 14 districts. Due to the scarcity of honey samples collected from some districts, it was not possible to work out all the properties of honey equally for all districts.

### 2.2. Honey harvesting and collection

A total of thirty nine (39) honey samples were collected by carefully excavating into the underground nest until reaching the nest chamber containing both honey and pollen stores. Fresh and pure honey samples of stingless bee (*Meliponula beccarii)* honey were harvested directly from sealed honey pots with disposable syringes. The collected samples were placed in plastic containers, packed in thermal boxes with ice, and taken to the laboratory (Holeta Bee Research Laboratory and other respected laboratories) and were further strained for impurities and stored in the refrigerator (−4°C) until the laboratory analysis was conducted. The physicochemical and microbiological analyzes were performed after the collection of samples not exceeding one month.

### 2.3. Analyzing the botanical origin of honey

Honey pollen analysis was done using the methods, recommended by Louveaux *et al*. [25]. For this purpose, 10 grams of honey was dissolved in 20 ml of warm distilled water and the sediment was concentrated by repeated centrifuging at 3800 rpm for 10 minutes and the supernatant was decanted. The distilled water of 20 ml again was added to completely dissolve the remaining sugar crystals and centrifuged at 3800 rpm again for 5 minutes and the supernatant was removed completely. The sediment was spread evenly using a sterile micro spatula on a microscope slide and the sample was dried for a while. Thereafter, one drop of glycerin jelly was added to the coverslip and the pollen grains were identified using a pollen atlas [26]. The percentage of pollen types in each honey sample was calculated based on the total number of different types of pollen grains counted in each sample. If pollen grains counted were greater than 45 %, used as predominant pollen (monofloral honey), while honey with no predominant pollen was used as multifloral honey [25]. The pollen count was done under the light microscope (Swift instrument international, Japan, high power 40x) linked to a computer.

### 2.4. Determination of the physicochemical analysis of honey

#### 2.4.1. Color analysis

The colours of the honey samples were measured using a Pfund colour grader. Approximately 100 g of honey was poured into the sample holder of the Pfund grader. Determination was based on the matching of the honey sample colours with the colour indexes present in the glass Pfund grader.

#### 2.4.2. Moisture content

The moisture content of the honey samples was determined using an Abbe refractometer (ABBE-5 Bellingham Stanley. Ltd, United Kingdom) that can be thermostated at 20°C and regularly calibrated with distilled water. Honey samples were homogenized and placed in a water bath until all the sugar crystals were dissolved. After homogenization, the surface of the prism of the refractometer was covered with honey and after 2 minutes refractive index for moisture was determined. The value of the refractive index of the honey sample was determined using a standard table designed for this purpose [27].

#### 2.4.3. pH and free acidity

From each honey sample, ten grams of honey was dissolved in 75 ml of distilled water in 250 ml beaker and stirred using a magnetic stirrer. The electrode of the pH meter (METTLER TOLEDO, CHINA) was immersed in the solution and the pH of honey was recorded. For measurement of free acidity, the solution was further titrated with 0.1 M sodium hydroxide (NaOH) solution to pH 8.30. For precision, the reading to the nearest 0.2 ml was recorded using a 10 ml burette. Free acidity is expressed as mill equivalents or a mill mole of acid/kg honey and is equal to ml of 0.1M NaOH × 10. The result is expressed to one decimal place following the procedure of the International Honey Commission [27].

Acidity =10 V, Where: V = the volume of 0.1N NaOH in 10 g of honey.

#### 2.4.4. Ash content

Determination of ash content was carried out by incinerating honey samples at 600°C in a muffle furnace (BioBase JKKZ.5.12GJ, Shandong, China) to constant mass [27]. First, the ash crucible was heated in an electrical muffle furnace at ashing temperature and subsequently cooled in a desiccator to room temperature and weighted to 0.001 g (M_2_). Then 5 g (M_0)_ of each honey sample was weighed to the nearest 0.001 g and taken into a platinum crucible and two or three drops of olive oil were added to prevent foaming. Water was removed and started ashing without loss at a low heat rise to 350 - 400°C using a hot plate. After the preliminary ashing, the crucible was placed in the preheated furnace and heated for at least 4-6h. The ash crucible was cooled in the desiccators and weighed. The ashing procedure was continued until a constant weight was reached (M_1_). Lastly, % of the weight of ash in g/100 g honey was calculated using the following formula: -

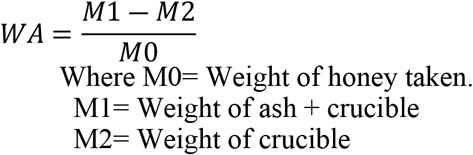

#### 2.4.5. Sugars profile

Honey sugars were determined using high-performance liquid chromatography (HPLC-1260 Infinity Series Agilent Technologies, Germany). Five grams of honey were dissolved in 40 ml of distilled water. A 25 ml of acetonitrile was pipetted into a 100 ml volumetric flask. The honey solution was transferred to a flask and filled to the mark with distilled water. Finally, the solution of each honey sample was filtered using a syringe filter (0.45 µm) before chromatographic analysis. The HPLC separation system was composed of an analytical stainless steel column, 4.6 mm in diameter, 250 mm in length, containing amine-modified silica gel with 5-7 µm particle size. Flow rate 1.3 ml/min, mobile phase Acetonitrile: water (80:20, v/v) and sample injection volume 10 µl. The sugars were detected by a Refractive Index Detector thermostated at 30°C temperature regulated column oven at 30°C. The identification of honey sugars was obtained by comparison of their retention times with those of the standard sugars [27]. Standard sugars with their percent purity level used were glucose (> 99.5%), sucrose ((> 90%), maltose (> 90%), and fructose (>99.5%) made in Germany, Sigma Aldrich. Five series serial dilution standards of fructose, glucose, sucrose, and maltose mixture which contain 2 g, 1.5 g. 1 g, 0.5 g, and 0.15 g were weighted and dissolved in 40 ml distilled water and 25 ml acetonitrile according to the International honey commission [27] to draw a calibration curve.

#### 2.4.6. Electrical conductivity

The electrical conductivity of a solution of 20 g dry matter of honey in 100 milliliters of distilled water was measured using an electrical conductivity cell (BANTE Instrument-520 conductive and temperature meter, China). A 0.745 g of potassium chloride (KCl), was dried at 130°C, dissolved in freshly distilled water in a 100 ml flask, and filled to volume with distilled water. Forty milliliters of the potassium chloride solution were transferred to a beaker and the conductivity cell was connected to the conductivity meter, the cell was rinsed thoroughly with potassium chloride solution and immersed the cell in the solution, together with a thermometer and reading of the electrical conductance of the solution in mill siemen after the temperature has equilibrated to 20°C was taken as described in harmonized international commission [27].

The cell constant K was calculated using the following formula:

K=11.691x1/G

Where:

K= the cell constant in cm^-1^

G= the electrical conductance in mS, measured with the conductivity cell. 11.691= the sum of the mean value of the electrical conductivity of freshly distilled water in mS.cm^-1^ and the electrical conductivity of a 0.1M potassium chloride solution, at 20°C.

#### 2.4.7. Hydroxyl methyl furfural (HMF)

HMF was determined using 6800 UV– Vis spectrophotometer (JENWAY, United Kingdom). A 5 g of honey sample was weighed in a small beaker and mixed in 25 ml distilled water and transferred into 50 ml volumetric flask [27]. A 0.5 ml carrezz solution I (15 g K_4_Fe (CN) _6_. 3H_2_0 /100 ml distilled water) was added and mixed with 0.5 ml carrezz solution II (30 g Zn acetate /100 ml distilled water). The solution was mixed with the honey solution. A droplet of alcohol was added to the solution. The solution was filtered through a filter paper and the filtrate (10 ml) was discarded. A 5 ml filtrate was added to each of the two test tubes and 5 ml distilled water was added to the first test tube (sample solution), while 5 ml sodium bisulfite solution (0.20% of 0.20 g NaHSO_3_/100 ml distilled water) was added into the other test tube (reference). The contents of both test tubes were well mixed by vortex mixer and their absorbance was recorded spectrophotometrically by subtracting the absorbance measured at 284 nm for HMF in the honey sample solution against the absorbance of reference (the same honey solution treated with sodium bisulfite, 0.2%) at 336 nm and the result was calculated and expressed according to international honey commission [27].

Hydroxyl methyl furfural (HMF)/100 g honey = (A_284_ - A_336_) *149.7 * 5*D/W sample Where A_284_= absorbance at 284, A_336 =_ absorbance at 336, 149.7= constant, 5= theoretical nominal sample weight and W= mass of honey sample, D=dilution factor where necessary

### 2.5. Determination of microbiological quality of honey samples

A 20 g of each honey sample was mixed with 180 ml sterilized distilled water and homogenized using an orbital shaker (110 rpm) for five to ten minutes and used as a stock solution for further serial dilution. A 0.1 ml from 10^−1^ serial dilution was pipetted onto the center of sterilized respective plates. Accordingly, 20-25 ml sterilized respective media was poured onto the plate containing the aliquots and well mixed. The inoculated plates were incubated at the appropriate temperatures for 24-96 h. The colonies were enumerated from the countable plates.

#### 2.5.1. Aerobic mesophilic counts

A 1ml of aliquots was taken from the stock and serially diluted in 9ml sterilized normal saline (0.85% NaCl). A 0.1 ml was taken from 1:10 dilution and poured in duplicate onto plates of sterilized plate count agar and incubated at 32°C for 48 hrs. Viable cells were counted using a colony counter and the result was recorded as CFU/ml of honey [28].

#### 2.5.2. Staphylococci counts

A 1ml of aliquots was taken from the stock and serially diluted in 9 ml sterilized normal saline (0.85% NaCl). A 0.1ml was taken from 1:10 dilution and poured in duplicate onto plates of sterilized Mannitol Salt Agar and incubated at 32°C for 36 hrs. After incubation, yellow colonies with yellow zones and colorless or red colonies with red zones were counted as staphylococci. Colony count was conducted using a colony counter and the result was recorded as CFU/ml of honey [28].

#### 2.5.3. Aerobic bacterial spore formers

For aerobic bacterial spore counts, 0.1 ml of the serial dilutions was heated in a water bath at 80°C for 10 min to kill vegetative cells and cooled rapidly in tap water. A 0.1 ml of aliquot was poured onto plates of Nutrient Agar (Oxoid). The grown colonies were counted as aerobic spore former bacteria after incubation at 30-32°C for 48 hrs. nutrient agar [28].

#### 2.5.4. Yeasts and molds

From appropriate serial dilution, 0.1 ml of aliquot was poured onto plates of Potato Dextrose Agar (Oxoid) containing (g/l) potatoes infusion, 200.00 g; dextrose, 20.00 g; agar, 15.00 g; chloramphenicol, 0.1 g; distilled water, 1000 ml with a pH value of 5.6±0.2 and incubated at 25-28 °C for three to five days. Smooth (non-hairy) colonies without extension and hairy colonies with extension at the periphery were counted as yeasts and molds, respectively [28].

### 2.6. Statistical analysis

The data obtained in the study were analyzed using One-way ANOVA and the data was expressed as mean and standard errors (±). For all the computations, SPSS version-20 statistical software was employed and the differences between mean values were significant at values of p < 0.05.

## 3. Results and discussions

### 3.1. The botanical origin of honey

The botanical origin analysis of the honey samples showed that the stingless honey were, mainly originated from twelve (12) different nectar source plant species including: *Schefflera abyssinica, Croton macrostachyus, Coffee arabica, Vernonia amygdalina, Guizotia scabra, Eucalyptus spp, Syzygium guineense, Pterolobium spp, Ilexmitis spp, Tephrosia vogelii spp, Plantago lanceolata and Trifolium rueppellianum* (Table 1). These plant species are also known as major and minor honey plants for *A. mellifera* bee’s honey in the areas.

**Table 1.** Relative frequency of nectar source plant species in each sample of the study areas (% distribution)

| Honey source (Districts) | *Schefflera abyssinica* | *Croton macrostachy* | *Coffea arabica* | *Vernonia amygdalina* | *Guizotia scabra* | *Eucalyptus spp* | *Syzygium guineense* | *Pterolobium* | *Ilexmitis* | *Tephrosia vogelii* | *Trifolium rueppellianu* | *Plantago lanceolata* |
| --- | --- | --- | --- | --- | --- | --- | --- | --- | --- | --- | --- | --- |
| Alle (N=4) | 6.22 | 0.34 | 0.92 | 20 | **50.82** | 20.60 |  | 1.10 |  |  |  |  |
| Didu (N=2) | 42.21 | 28.69 | 2.86 | 3.43 | 4.74 | 14.38 | 3.43 |  |  |  |  |  |
| Gechi | **61.7** | 11.9 | 2.6 |  | 1.2 | 6.2 | 16.4 |  |  |  |  |  |
| Gera (N=2) | 41.38 | 14.96 |  | 9.94 | 20.65 | 13.08 |  |  |  |  |  |  |
| A/J/M/B (N=3) |  |  | 18.18 | 13.16 | 10.91 | 23.12 | 34.63 |  |  |  |  |  |
| A/Sorraa (N=4) |  |  | 15.23 | 5.20 | 1.42 | 21.98 | 17.33 |  | 38.84 |  |  |  |
| Chalia (N=3) |  |  |  |  | 2.1 | **79.67** |  |  |  | 18.32 |  |  |
| Guduru |  |  |  |  | **73.2** | 13.0 |  |  |  |  | 2.39 | 11.41 |
| J/Ganat |  |  |  |  | **93.8** |  |  |  |  |  |  | 6.2 |
| Diga |  |  |  |  | **92.1** |  |  |  |  |  | 7.9 |  |
| Sibu Sire |  |  |  | 2.4 | **97.6** |  |  |  |  |  |  |  |

A/J/M/B=Arda Jilaa Me'ee Bokkuu; J/Ganat=Jimma Ganat; A/Sorra=Annaa sorraa, N= number of sample

The relative pollen frequency of honey samples indicated that *Schefflera abyssinica* was a predominant (61.7% of the pollen grains counted) honey plant in honey samples collected from the Gechi district of Buno Bedelle. In addition, *Schefflera abyssinica* was a secondary pollen source (41.38 %) in the Gera district of Jimma Zones and Didu (42.38%) district of I/A/Bora. *Schefflera abyssinica* is an abundant bee forage plant in moist highlands of southwestern and southeastern parts of Ethiopia and it provides monofloral honey in these areas[29]. Degaga[30] also reported that *Schefflera abyssinica* is a major monofloral honey source tree in Gera District, Jimma Zone.

*Guizotia* species was predominant in honey samples collected from Guduru and Jimma Ganat districts of Horo Guduru Wollega, Diga, and Sibu sire districts of East Wollega and Alle districts of I/A/Bora zones. A similar finding was also reported by Arega *et al*.[31] that the *Guizota* species is the major honey source plant that provides monofloral honey in Sibu sire and J/Ganat East Wollega and Horro Guduru Wollega respectively. The majority of bee plants flower after the heavy rainy season in July through September and most of the Ethiopian highlands are covered with golden-yellow flowers of *Guizotia spp*. and *Trifolium spp*. with many different colors[32]. While, *Eucalyptus* species were predominant in honey samples collected from the Chalia districts of West Shoa Zone and a secondary honey plant for the honey sample collected from A/J Me’ee Bokku, and A/Sorra districts of Guji Zone. In addition, *Coffee arabica* and *S. guineense* were also secondary pollen (16-45 %) for the honey sample collected from Guji Zone (A/J/M/Bokku and A/Sorra districts) (Table 1). *C. arabica* is a common cash crop in western Oromia and contributed much to monofloral honey samples. Adi *et al*. [33] stated that *C. arabica* flowers provide abundant pollen and nectar in January for honeybees. The study conducted by Bereke *et al*.[34] indicates that *C. arabica* has the potential to produce 125 kg/ha of honey from coffee plants. *S. guineense* is foraged by bees vigorously for the abundant nectar and pollen from the flowers and this tree is an important honey source for the country[32]. Similarly, a study conducted by Bareke and Addi[35] also indicated *Syzygium guineense* is the predominant plant species in the A/Sorra district of the Guji Zone from March to April.

*Ilexmitis* was a secondary honey plant in the honey sample collected from Guji of A/Sorra District. Bareke and Addi [35] also mentioned that *Ilexmitis* and *Syzygium guineense* were the major honey source plants that provide monofloral honey in this district. In addition to predominant frequency, different levels of abundance of given pollen type in nectar such as secondary, tertiary, and quaternary enrichment are required for the botanical description of honey[36]. The predominant pollen, secondary pollen, important minor pollen, and minor pollen species in enriching the stingless bee honey samples from the study area were presented in Table 1.

The categories of pollen sources from the honey samples from Table 1 above can be seen that predominant pollen source (> 45%), secondary pollen source (16-45%), important minor pollen source (3-15%), and minor pollen source (< 3%)

### 3.2. Physicochemical properties of different sources of stingless bee honey

The analysis of colour in our honey revealed Pfund scale ranged from 0 to 114 mm, categorized as water white to dark amber (Table 2). Stingless bee honey samples collected from A/J/M/Bokku and A/Sorra districts were found to be extra light amber, whereas the sample collected from Chalia, Jimma Ganat, Sibu sire, Diigaa, Didu, and Gera districts were classified as light amber. The *M*.*beccarii* honey collected from Guduru, Sokorru, and Seka was found to be amber and a sample collected from the Gechi district was the only sample grouped under water white. According to Color Standards Designations for extracted Honey [37]; the color of the *M*.*beccarii* honey for the present study was classified as 50 % light amber, 21.5% extra light amber, and amber, and 7 % water white. This describes that stingless bee honey comprises different colour. This may be attributed to different factors such as a method of production, light exposure, heat, and storage time, as well as enzymatic reactions. In most cases, the choice of, colour of the honey is based on consumers’ preferences which are considered as quality appreciation and market acceptability [38].

**Table 2.** Color of honey samples grouped according to the Pfund scale

| Honey samples source | Color name | Pfund scale (mm) |
| --- | --- | --- |
| Gechi | Water white | 0-8 |
|  | Extra White | >8-17 |
|  | White | >17-34 |
| Alle,A/J/M/Bokku,A/Sorra | Extra Light Amber | >34-50 |
| Chelia,J/Ganat,Sibusire,L/Assabii,Diigaa,D<br>idu,Gera | Light Amber | >50-85 |
| Gudurru,Sokorru,Seka | Amber | >85-114 |
|  | Dark Amber | >114 |

In the present study, the highest moisture content was recorded for the honey sample collected from the Sibu Sire district (38.3±0.10%) and Sekka district (38.1±0.58%) while the lowest was recorded for stingless bee honey collected from L/Assabi (25.17±0.18%) (Table 3). The moisture contents of the present study agreed with Biluca *et al*. [39] who indicated that the moisture content of stingless bee honey ranged from 23.1 to 43.5g/100g in Brazil and Chuttong *et al*.[11] reported 25 to 47 g/100 g for Thailand. Souza *et al*. [40] also mentioned the variations of moisture content ranged from 19.90 % to 41.90 % in the honey of different species of stingless bees. The moisture contents of honey samples in the present study (24.50–39.0%) were almost similar to the previously reported moisture content values of stingless bee honey produced in the West Shoa zone 25 to 35% [22].

**Table 3.**
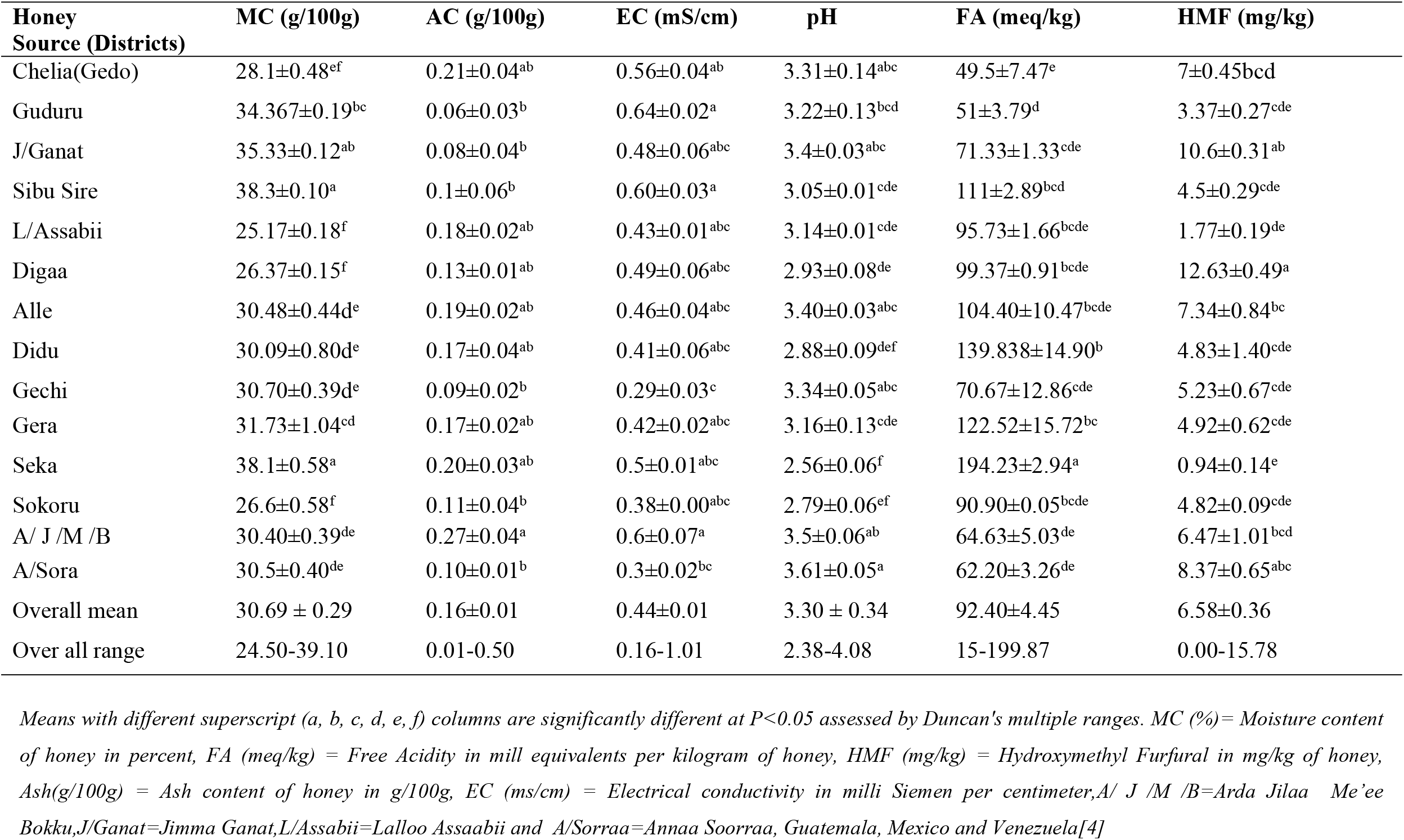
The mean and standard error (mean ± SE) values for the physicochemical content of stingless honey collected from study areas.

The ash content of the samples collected from A/ J/ M/Bokku was significantly different (P<0.05) from the honey samples collected from A/Sorra, Sokoru, Sibu Sire, J/Ganat, and Guduru districts. The highest mean ash content was recorded for stingless bee honey collected from A/J/M/Bokku (0.27±0.04 g/100 g) and the lowest was recorded for the honey sample from Guduru(0.06±0.01 g/100 g) (Table 3). The overall range of ash content (0.01-0.50g/100g) of the present study was comparable with the previous finding (0.41%) in West Shoa, Ethiopia,[22]. The present result was also consistent with Suntiparapop *et al*. [41]work that deals with ash content for Tetragonula laeviceps of Thailand, with 0.20 g/100g. Gonzalez *et al*.[42]also mentioned 0.01 to 0.6% with a mean of 0.16 ± 0.12% of ash value for the honey samples analyzed in Mexico. However, Chuttong *et al*. [11] presented higher ash content of 0.22 to 3.1 g/100 g for honey from stingless bee species from Thailand.

The electrical conductivity of *M. beccarii* of honey samples collected from Guduru (0.64 ± 0.02 mS/cm) was significantly different (p < 0.05) from the EC values of the honey sample from the Gechi and A/Sorra. The electrical conductivity in the current study ranged from (0.16-1.01mS/cm) with a mean value of 0.44 ± 0.02mS/cm is in line with those of the previous study (0.22-1.52mS/cm) stingless bee honey in Ethiopia [22], (0.58 ± 0.14) reported by Alvarez-Suarez *et al*. [43] for *Meliponula beccarii* honey in Cuban, (3.27±0.01) for *Trigona* species honey in Ethiopia [44]. Suntiparapop *et al*.[41] also reported electrical conductivity of 0.71 and 0.53 ms/cm from South East Asia. Similar findings were also mentioned by Sousa *et al*. [45] and Guerrini *et al*.[46], pointing to values of 0.30 and 0.67 mS cm^-1^ respectively from south America. The finding of the current study was lower when compared to the electrical conductivity of honey from stingless bee spices in Brazil ranging from 0.15 to 1.34 mS/cm [47]. In general, variations in electrical conductivity of honey samples were linked to variations in the geographical, botanical, and entomological origins of the honey samples [39].

The overall pH values of the *M. beccarii* of the honey sample obtained from this study range (from 2.38 to 4.08) with a mean value of 3.30. The highest mean pH value (3.61± 0.05) was recorded for stingless bee honey from the A/ Sorra district of the Guji Zone and the lowest (2.56± 0.06) was recorded for the honey sample in the Seka District of Jimma Zone. This is consistent with recent studies indicating that pH values are 3.2 to 4.5 in West Shoa of Oromia[22] and 3.2±0.21 in Brazil [48]. Overall, the honey of a stingless bee has been characterized by low pH, because of the presence of different acids (gluconic, acetic, lactic, citric, succinic, formic, malic, malic, and oxalic), and a reduced total solute content [49]. This low pH of honey may contribute to the antibacterial activity as it is believed to prevent the growth of bacteria [50].

The free acidity value varied from 15 to 199.87 meq/kg with a mean value of 92.29 ±4.45 meq/kg. The free acidity of the Seka District of the Jimma zone (194.23 ± 2.94 meq/kg) was significantly (p < 0.05) different from all sources of honey samples. The lowest mean value of free acidity 49.5± 7.47 meq/kg was recorded for the honey sample from Chalia of the West Shoa Zone. The free acidity of *M. beccarii* from this study was comparable with 136.8±7.6 meq/kg for Malaysia stingless bee [51] 124.2 ± 22.9 meq/kg for Australia stingless bee [52], *Melipona quadrifasciata* and *Tetragonisca angustula* from Brazil with average 103.3 meq/kg and 109.0 meq/kg [53], respectively.

Variations in free acidity may be related to the harvest season, the maturity of honey, floral sources, locations, storage condition, and/or climatic factors, which would favor chemical, enzymatic, and microbiological reactions able to release acidic compounds in honey [54,55]. Following the known high moisture content of stingless bee honey, free acidity values were also frequently reported to be higher in stingless bee honey compared to the *Apis mellifera* honey[53,56]. Free acidity indicates one of the quality parameters of honey samples and it reveals whether the honey is fermented or not [57]. International regulations specify that free acidity is not higher than 50 meq/kg of honey for *Apis mellifera* [5,24]. For this study, except for the honey sample from the Chalia Districts, the mean free acidity of other honey samples doesn’t fit the international [5,24].

The result of this finding indicated that the highest concentration of HMF (12.63 ± 0.49 mg/kg) was recorded for *M. beccarii* honey collected from the Diiga district of the East Wollega Zone. The HMF value of the current study varied from 0.00 to 15.78 mg/kg with a mean value of 6.58 ± 0.37 mg/kg. These findings are in agreement with those obtained by Oddo *et al*. [52], who mentioned the HMF contents ranging from 0.4 to 2.1 mg/kg, and Suntiparapap *et al*.[41] who reported the HMF value of 0.53 to 0.71 mg/kg of honey for stingless bee honey collected from Australia and Thailand, respectively. However, the average HMF value of the current study (6.58 ±0.67 mg/kg) is lower than the previous study conducted by Gela *et al*. [22] reported 18 mg/kg in the West Shoa zone of Oromia. The HMF value of the investigated samples is within the range of the established national [23] and international [5,24] quality standards that permit a maximum of 40 mg/kg. This amount of HMF in honey is one of the important indicators of honey’s quality (freshness) indicating whether the honey is aged or over-heated[58,59].

#### Sugar profile of stingless bee honey

In this study result, the highest fructose content was recorded for the honey sample collected from Gera (40.29±0.72%) and the lowest was recorded for Sokorru (18.27±0.37%) of Jimma Zone (Table 4). The overall fructose content ranged between 16.53-47.50 g/100 g with a mean value of 36.48 ± 0.54 g/100 g, which agrees with the finding by Almeida-Muradian *et al*. [60] indicating 31.61 g/100 g for the stingless bee species of *M. beccarii* in the Amazon region of Brazil. The concentration of glucose for honey samples collected from the Gechi district was significantly highest (36.2±2.33g/100g) and a honey sample collected from the Seka district (20.76 ±0.69 g/100 g) was recorded for the lowest glucose level. From this finding, the glucose content ranged from 16.88 to 40.24 g/100g with a mean value of 27.67 g/100 g (Table 4) which is in line with those of previous studies by Almeida-Muradian *et al*. [60] indicating 33 g/100 g for species of Melipona in the Amazon region of Brazil. However, the glucose level of the current study was higher than those reported in previous studies by Guerrini *et al*.[46] indicating 25.5 g/100g for stingless bee honey in Ecuador, Rizélio *et al*.[61].

**Table 4.** The mean and standard error (mean± SE) of the sugar profile of stingless bee honey

| Honey Source (Districts) | Fructose (%) | Glucose (%) | Sucrose (%) | Maltose (%) | RS (%) |
| --- | --- | --- | --- | --- | --- |
| Chalia | 34.31±0.27<sup>ab</sup> | 25.88±0.69<sup>bcd</sup> | 1.22±0.44<sup>b</sup> | 0.51±0.05<sup>b</sup> | 60.19±0.88<sup>cd</sup> |
| Guduru | 38.83±0.08<sup>ab</sup> | 25.49±0.35<sup>bcd</sup> | 2.82±0.08<sup>ab</sup> | 0.25±0.01<sup>b</sup> | 64.32±0.27<sup>bcd</sup> |
| J/Ganat | 39.48±0.70<sup>ab</sup> | 26.56±0.16<sup>bcd</sup> | 0.07±0.00<sup>b</sup> | 0.6±0.01<sup>b</sup> | 66.04±0.86<sup>bcd</sup> |
| Sibu Sire | 40.02±0.59<sup>a</sup> | 28.23±0.48<sup>b</sup> | 0.18±0.02<sup>b</sup> | 0.51±0.00<sup>b</sup> | 68.24±0.11<sup>abc</sup> |
| L/Assabii | 37.38±0.04<sup>ab</sup> | 27.65±0.53<sup>bc</sup> | 0.26±0.01<sup>b</sup> | 0.68±0.02<sup>b</sup> | 65.03±0.57<sup>bcd</sup> |
| Diigaa | 36.67±0.15<sup>ab</sup> | 28.31±0.03<sup>b</sup> | 1.56±0.11<sup>ab</sup> | 0.43±0.01<sup>b</sup> | 64.97±0.13<sup>bcd</sup> |
| Alle | 32.54±1.52<sup>b</sup> | 28.57±1.16<sup>b</sup> | 0.75±0.20<sup>b</sup> | 1.25±0.17<sup>b</sup> | 61.11±1.81<sup>cd</sup> |
| Didu | 38.75±0.63<sup>ab</sup> | 24.93±1.11<sup>bcd</sup> | 2.87±0.84<sup>ab</sup> | 2.70±1.08<sup>ab</sup> | 63.68±1.17<sup>bcd</sup> |
| Gechi | 39.93±1.31<sup>a</sup> | 36.2±2.33<sup>a</sup> | 0.17±0.01<sup>b</sup> | 0.89±0.00<sup>b</sup> | 76.13±3.64<sup>a</sup> |
| Gera | 40.29±0.72<sup>a</sup> | 31.25±2.62<sup>ab</sup> | 0.37±0.11<sup>b</sup> | 1.08±0.24<sup>b</sup> | 71.5±3.15<sup>ab</sup> |
| Seka | 36.72±0.36<sup>ab</sup> | 20.76±0.69<sup>d</sup> | 0.36±0.18<sup>b</sup> | 4.13±0.36<sup>a</sup> | 57.48±0.44<sup>d</sup> |
| Sokorru | 18.27±0.37<sup>c</sup> | 21.63±0.91<sup>cd</sup> | 4.01±0.50<sup>a</sup> | 0.32±0.20<sup>b</sup> | 39.9±1.10<sup>e</sup> |
| A/ J/ M /B | 38.61±0.96<sup>ab</sup> | 27.38±0.73<sup>bc</sup> | 1.12±0.40<sup>b</sup> | 0.53±0.06<sup>b</sup> | 65.99±1.46<sup>bcd</sup> |
| A /Sooraa | 37.50±0.94<sup>ab</sup> | 28.43±0.47<sup>b</sup> | 1.18±0.41<sup>b</sup> | 0.99±0.39<sup>b</sup> | 65.93±1.20<sup>bcd</sup> |
| Overall mean | 36.48± 0.54 | 27.67±0.43 | 1.24±0.17 | 1.19±0.18 | 64.19±0.75 |
| Overall Range | 16.53-47.50 | 16.88-40.24 | 0.0-9.0 | 0.0-12 | 36.32-72.44 |
| Vit *et al.*, 2004<br>(*Meliponinae*) | - | - | Max 6 | - | Min 50 |
| Ethiopia *Apis mellifera* | - | - | ≤ 10 | - | ≥65 |

Note: Means with different superscripts (a, b, c, d, e) within the columns are statistically different at $P < 0.05$ . RS (%) = percent of Reducing Sugar in honey, Ethiopian Standards Agency[23] and = Guatemala, Mexico, and Venezuela [4]

**Table 5.**
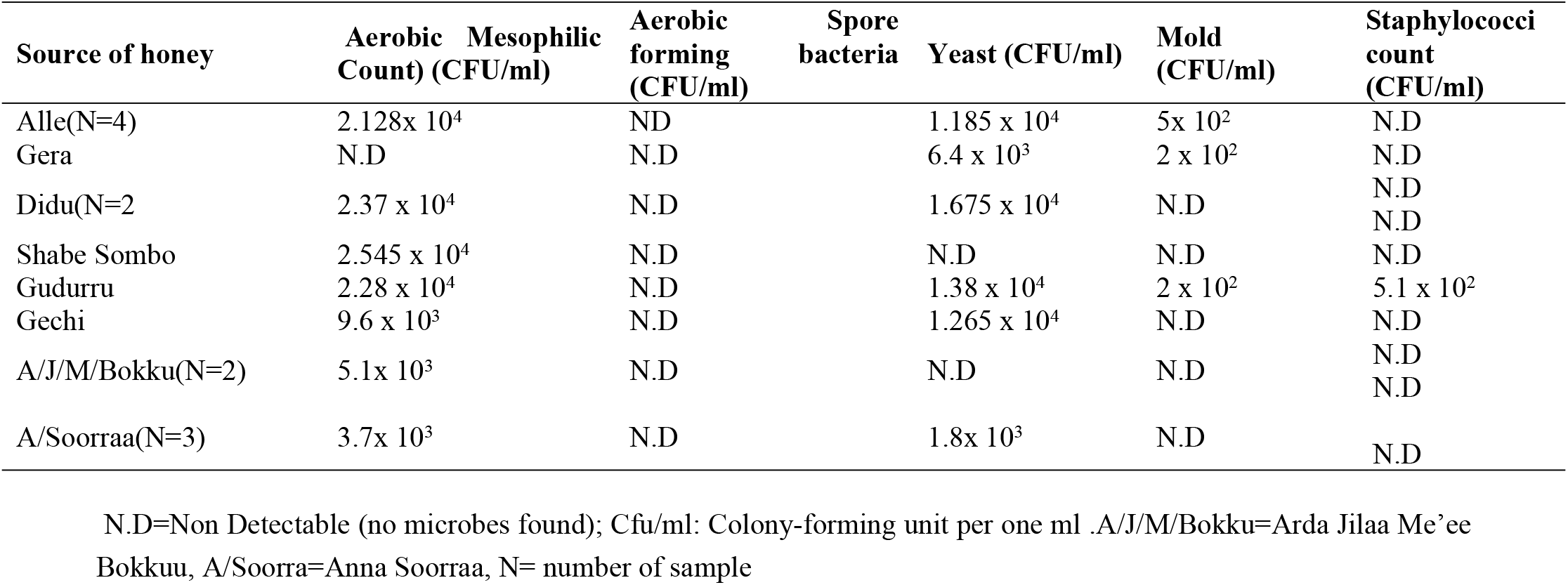
Microorganisms isolated from stingless honey samples from the study districts

The results of the present study support the hypothesis that fructose was more abundant than glucose dominant sugars in a good quality stingless bee honey may be responsible for the intensity of the sweet taste and high hygroscopicity while maintaining the liquid honey for a long time or never crystallize.

The mean value of the sucrose content of *M. beccarii honey sample* collected from Sokoruu was a statistically significant difference (p<0.05) among all honey except honey samples collected from Didu, Guduru, and Diiga districts. The highest concentration of sucrose was recorded for honey samples collected from the Sokkoru district of the Jimma Zone (4.01±0.50) while the lowest was recorded for the J/Ganat district of the H/G/Wollega Zone (0.07±0.00) (Table 4). The present results are consistent with Souza *et al*.[53] work that indicated the sucrose ranged from 0.13 to 6.00 g/100 in stingless bee honey from Brazil. As for indicators of quality, sugars are related to honey maturation. High levels of non-reducing disaccharides or sucrose that have not yet been converted to glucose and fructose may be indicative of premature harvesting of the product [56]. All of the stingless bee honey samples fit the standard of national [23]and international[5,24] quality requirements with a maximum of 10% and 5% sucrose content from *Apis mellifera*, respectively. The quality standard of stingless bee honey was not legislated and a comparison was carried out with the norm by[4], who proposed a maximum of 6% for the *M. beccarii* honey. This agrees with the sucrose content of the finding obtained in this study. These results demonstrate that honey was collected at the ideal maturation time because high sucrose content may result from the addition of commercial sugar or may be attributed to the early honey collection.

The maximum and minimum average of maltose was recorded from the Seka District of Jimma Zone (4.13 ± 0.36 %) and Guduru district (0.25±0.01%), respectively (Table 4). From the present analysis, the Seka honey sample was a statistically significant difference (P<0.05) from other honey samples except the honey sample collected from the Didu district of I/A/Bor Zone. A similar finding was reported by Vit *et al*. [49] suggested that the genus *Meliponula beccarii* produces honey with low maltose content (1-1.23 g/100 g), whereas the genus *Trigona*, such as *T. angustula*, produces honey with high maltose content (24-56 g/100 g). Kaškonienė [62] suggested that artificial honey was distinguished from natural honey by a high amount of maltose.

In this present study, the reducing sugar values ranged from 36.32 to 72.44% with a mean value of 64.15 ± 0.75%. These results are consistent with the finding by Carvalho *et al*. [63] who indicated the variations from 42.55 % to 55.61%, and Souza *et al*. [40] reported the value ranging from 58.0 to 75.7%. Alves *et al*.[64] also mentioned the levels of reducing sugars in *Meliponula beccarii* from 50.60% to 95.60%. However, the study by Chuttong *et al*. [9] indicated an average lower than 60% for reducing sugar content. These contrary findings suggest that the sugar profile of honey from stingless bees can vary from one region to another, depending on the flora and vegetation that predominates in that region [65]. All the samples of the current study atctained the minimum amount of reducing sugars (50 g/100 g) established by Vit *et al*. [4] as the standard of quality for Melipona honey. However, except for honey sample collected from Guji (A/Sorraa &A/J/M/Bokku), the reducing sugar of *M. beccarii* from Gera, Gechi, L/Assabii, J/ganat, and Sibu Sire districts were much less and did not fit the standard international quality requirements[5,24] which should be a minimum of 65% for *A. Mellifera*. These results indicate that the sample of stingless bee honey might not be harvested during its appropriate time or not matured, which can alter the normal values of carbohydrate composition in the honey. Evidence has shown that stingless bee honey contains lower sugar content, is less sweeter, and has higher moisture compared to *Apis mellifera* [11,45,47].

#### Microbiological profiles of honey samples

Microbiological contamination of honey can be caused by pollen microbiota, from the bee itself, or by failures in the hygiene of the manipulator during the processing of the product [71]. According to our study, the highest Aerobic Mesophilic (2.545 × 10^4^ CFU/ml) and least mold (2 × 10^2^ CFU/ml) were counted from the stingless bee honey sample collected from Shabe sombo of Jimma Zone and Gera of Jimma Zone as well as Guduru district of Horo Guduru Wollega Zone respectively. From this finding, the majority of the stingless bee honey samples were positive for Aerobic Mesophilic. The *M. beccarii* honey sample collected from Shabe sombo of Jimma Zone had the highest 2.545 × 10^4^cfu/ml and a sample collected from A/ Sorraa of Guji Zone had the lowest mean count of 1.9 × 10^3^ CFU/ml of Aerobic Mesophilic. Relatively as compared with other microbes, a high population of Aerobic Mesophilic Count was recorded which could be an indicator of poor handling and contact with the bare hands of honey handlers.

The presence of the aerobic microorganisms is an indicator of the degree of deterioration in honey and can help determine the useful shelf-life of honey [66]. However, some characteristics of honey, such as acidity, low water activity, low protein content, and high sugar, can diminish or stop bacterial activity, contributing to the longer shelf-life of the product [24,67]. Some of the samples from this study showed a high count of mesophilic bacteria, maybe due to quality and safety problems that can result in contamination. These sources of contamination include soil, water, air, pollen, and nectar [68,69]. On the other hand, the number of spore-forming bacteria and Staphylococci were at non-detectable levels in all samples (except the sample from B-Guduru has a staphylococci count). The low number of staphylococci can be explained by the aseptic collection of honey, equipment, and physical honey handling are considered the main sources of contamination by this microorganism [70,71].

Regarding the yeast, the stingless honey (*M. beccarii)* collected from the Didu district of the I/A/Bor Zone had the highest mean count of 1.675 × 10^4^ CFU/ml, and 9 × 10^2^ CFU/ml is the lowest mean count of yeast recorded for the A/J/ M/Bokku district of Gujii zone. Pinheiro *et al*. [72] mentioned the average count of fungi and yeast was 9.12×10^3^CFU/g for stingless bee honey in Brazil. In addition, Camargo *et al*. [73] suggested a limit of 1.0×10^4^CFU/g for the honey from the stingless bee, as the elevated humidity of honey is associated with the rich fungi microbiota of stingless bee’s honey resulting in a product with higher counts. Thus, the main problem is caused by their presence in the fermentation of honey, which can reduce the shelf life of the product[74]. Bogdanov *et al*.[75] presented the relationship between the moisture content of honey and fermentation risk and stated that honey with a moisture content of 17.1% to 18% is safe from fermentation if the yeast count is below 1000 *CF U/g*. Honey with 18.1 to 19% moisture content is safe from fermentation if the yeast count is below 10 *CF U/g*.

The highest mold means counted of 1.8 × 10^3^ CFU/ml was recorded A/ Sorra district of Guji and the lowest mean counted of mold 2 × 10^2^CFU/ml was recorded for honey samples collected from Gudurru district of Horo Guduruu Wollega and Gera district of Jimma Zone. The presence of molds in this study might be due to unhygienic practices during harvesting, packaging, and storing of the honey samples. Molds are known as xerophiles since they thrive in samples with low water contents between 16.2 to 17.0%. The high water content of stingless bees’ honey could favor the growth of molds and yeasts [76]. The absence of yeasts and molds in some honey samples confirms that honey has inherent antimicrobial properties that can delay the growth of many microorganisms. Osmophilic microorganisms, such as molds and yeasts, present the greatest risk to the quality of the honey because they can survive in acidic conditions and are not inhibited by sugar [77]. The low counts and limited variety of microbes are expected because of honey’s antibacterial properties against the growth or persistence of many organisms [78,79]. It is believed that the amounts of microorganisms in honey are lower than in any other natural food due to their high sugar concentration [80]. In addition to this factor, the presence of phytochemical molecules such as phenols, terpenes, and pinocembrine helps control the growth of microorganisms in honey [81]. The presence of a certain amount of bacteria and fungi in the samples is considered to be associated with the contamination during harvesting, straining, transportation, and storage [82,83].

## Conclusion and Recommendation

In this present work, honey samples from different agro ecology of Oromia have been analyzed for their quality criteria concerning physicochemical and microbiological quality profiles. The majority of physicochemical properties of analyzed samples are within the maximum limit of the quality criteria set by the Ethiopian Standard and Codex Alimentarius for *Apis Mellifera* except for moisture content, free acidity, and sugar profile. This variation demonstrates the differences between the honey samples of *Apis mellifera* and *Meliponula beccarii* and the need for specific standard regulations for stingless bee honey. Although stingless bee honey is known for its low pH value and antibacterial properties; however, microbial growth has been found in most samples, expressing poor hygienic procedures during harvesting, handling, transportation, and storage. The presence of *Staphylococcus* species in the *Meliponula beccarii* honey sample from Guduru might be due to improper handling and hygiene during honey harvesting and processing since *Staphylococcus*is amongst the most common pathogens found on hands. On other hand, Aerobic mesophilic is the indicator of the quality and safety problems during honey harvesting. Some *Meliponula beccarii* honey samples are contaminated with yeast, mold, and bacterial organisms indicating inadequate hygiene conditions during harvesting, handling, processing, and/ or storage. According to our knowledge, this study is the first comprehensive physicochemical and microbiological properties of honey conducted on stingless bee species (*Meliponula beccarii*) in the country from broader agro-ecological zones of Oromia, Ethiopia. In general, the current legislation regarding *Apis mellifera* honey is not adequate for all the characteristics analyzed, emphasizing the need for a suitable standard for stingless bee honey to prevent adulterations and to allow its commercialization on a formal market. Indeed, improved methods of harvesting methods could be developed to reduce the introduction of microorganisms and the quality standard should be proposed for stingless bee honey.

## Authors’ contributions

Conceptualization: Teferi Damto.

Data curation: Teferi Damto, Dheressa Kebaba,Meseret Gemeda

Formal analysis: Teferi Damto, Dheressa Kebaba,Meseret Gemeda

Funding acquisition: Teferi Damto

Methodology: Teferi Damto, Dheressa Kebaba,Meseret Gemeda.

Software: Teferi Damto

Visualization: Teferi Damto, Dheressa Kebaba,Meseret Gemeda.

Writing – original draft: Teferi Damto

Writing – review & editing: Teferi Damto, Dheressa Kebaba, Meseret Gemeda

## Funding

This work was supported by Ethiopian Institute of Agricultural Research (EIAR).

## Data Availability Statement

Data included in article/supplementary material/referenced in article.

## Acknowledgments

The authors are thankful to Holeta Bee Research Center for logistic support for the entire activities. Our special thanks also go to technical staffs of Bee product quality improvement and value addition research team and participant experts from respective districts for their support during field activities and sample collection

## Conflicts of Interest

The authors declare that they have no known competing financial interests or personal relationships that could have appeared to influence the work reported in this paper.

## Notes

### Competing Interest Statement

The authors have declared that no competing interests exist.

